# Quantitative phase imaging with temporal kinetics predicts hematopoietic stem cell diversity

**DOI:** 10.1101/2024.09.29.615639

**Authors:** Takao Yogo, Yuichiro Iwamoto, Hans Jiro Becker, Takaharu Kimura, Ayano Sugiyama-Finnis, Tomomasa Yokomizo, Toshio Suda, Sadao Ota, Satoshi Yamazaki

## Abstract

Innovative identification technologies for hematopoietic stem cells (HSCs) have advanced the frontiers of stem cell biology. However, most analytical techniques capture only a single snapshot, disregarding the temporal context. A comprehensive understanding of the temporal heterogeneity of HSCs necessitates live-cell, real-time and non-invasive analysis. Here, we developed a prediction system for HSC diversity by integrating single-HSC ex vivo expansion technology with quantitative phase imaging (QPI)-driven machine learning. By analyzing single-cell kinetics with QPI, we discovered previously undetectable diversity among HSCs that snapshot analysis fails to capture. Our QPI-driven algorithm quantitatively evaluates the stemness of individual HSCs and incorporates temporal information to significantly improve prediction accuracy. This platform marks a paradigm shift from “identification” to “prediction”, enabling us to forecast HSC status by analyzing their past temporal kinetics.

## Main Text

Hematopoietic stem cell (HSC) research has advanced in parallel with the development of innovative identification technologies. Originally, the spleen colony-forming assay was employed to evaluate the differentiation capacity of HSCs (*1*). However, the advent of flow cytometry in the 1980s marked a paradigm shift in cell categorization methodology (*2*), unveiling the presence of murine HSCs in a subset defined as Thy-1^low^ Lineage^-^ Sca1^+^ (*3*). Subsequent investigations elucidated the existence of long-term self-renewing HSCs (LT-HSCs) and transient short-term HSCs (ST-HSCs) (*4*). Ultimately, LT-HSCs were directly identified at the single-cell level within a bone marrow fraction characterized as Lineage^-^ cKit^+^ Sca1^+^ CD34^-^ (*5*). Similarly, human HSCs have been identified within the CD34^+^ CD38^-^ CD90^+^ CD45RA^-^ CD49f^+^ fraction of human umbilical cord blood, also at single-cell resolution (*6*). In line with these discoveries, we have previously reported on the exploration of HSC-specific marker genes and the development of methods for marking HSCs both in vitro and in vivo, establishing a technical platform (*7*-*9*).

These advances in HSC identification techniques have driven the entire field of stem cell research forward, leading to the discovery of numerous stem cell-specific markers (*10*-*12*). Moreover, single-cell analytical methods have uncovered the complex heterogeneity within stem cell populations (*13*, *14*). Specifically, in the field of hematology, this heterogeneity can be observed in the differential differentiation capacity of HSCs. Some HSCs are predominantly committed to myeloid or lymphoid lineages, while others are biased towards megakaryocyte differentiation (*15*-*18*). Additionally, heterogeneity exists in the metabolic states and clonal expansion capacities of HSCs (*19*-*21*). Single-cell RNA sequencing (scRNAseq) and other technologies have emerged as established tools for elucidating the diversity among individual cells, significantly advancing research into the heterogeneity of HSCs (*22*). However, these techniques are limited to evaluating the cellular state at a single temporal snapshot, thus failing to capture temporal dynamics. Cells are dynamic system, they undergo continual change and never retain a single state, HSCs are no exception (*23*, *24*). The comprehensive understanding of the temporal diversity of HSCs mandates an experimental system that allows for the real-time tracking and non-invasive probing of live cells for their analysis. The ultimate objectives include forecasting future cellular status based on past temporal kinetics and quantitatively predicting the functionality of HSCs at the single-cell level. Such predictive systems will be pivotal in further advancing the field of stem cell biology through enabling the establishment of a novel platform that predicts future stem cell diversity using their past behavior.

The long-term evaluation of phenotypic and kinetic behaviors of single HSCs firstly requires a culture system that allows for HSC expansion while maintaining function. We have previously established a long-term culture system for the expansion of both murine and human HSCs (*25*-*27*). Conventional fluorescent imaging techniques may impair stem cell function due to the introduction of fluorophore or high-intensity illumination, potentially causing phototoxicity. Ptychographic quantitative phase imaging (QPI) techniques facilitate non-invasive and label-free monitoring of live cells across a wide field of view without the need for high-intensity light imaging (*28*, *29*). Furthermore, the meniscus compensation step during phase reconstruction actively senses the deformation of transmitted light, enabling fully quantitative and aberration-free imaging, even in U-bottomed culture wells, which allows high-throughput single HSC imaging during ex vivo expansion. In this work, by integrating our single-HSC ex vivo expansion technology (*25*, *26*) and QPI-driven machine learning, we have developed a prediction system for HSC diversity. This achievement signifies a paradigm shift from the era of “identification” of HSCs through snapshot analysis to the era of “prediction” of HSC function based on temporal kinetics, thereby fundamentally altering the landscape of stem cell research.

## Results

### Quantitative phase imaging and single-cell kinetics analysis reveal infinite diversity of HSCs

To investigate the kinetics of HSCs during ex vivo expansion at a single-cell level, we combined our recently established single-cell expansion culture system for murine HSCs with QPI. We sorted a single CD201^+^CD150^+^CD48^-^KSL cell from a population of ex vivo expanded HSCs into a 96-well U-bottom plate and monitored expansion for 96 hours with time-lapse QPI (Fig. 1A). Quantitative analysis of cellular kinetics revealed remarkable diversity. After 96 hours, marked differences in proliferation rate could be observed where 12.5% of HSCs underwent a rapid proliferation defined as producing more than 20 cells whereas other HSCs divided more slowly with 21.9% producing less than 4 cells (Fig. 1, B to F, Movie S1). Morphologically, we observed significant variations in the output cells; 10.9% of HSCs produced cells with dry masses larger than 200 pg whereas 17.2% of HSCs produced cells with dry masses smaller than 100 pg (Fig. 1, G to K, Fig. S1, A and B, Movie S2). We further investigated the interval between the first and second cell divisions (Division Gap) and observed significant variations among cells; in 25.5% of cells, this interval exceeded 5 hours, indicating the possibility of asymmetric cell divisions (Fig. S1C, Movie S3). Interestingly, we also observed interrupted cytokinesis during the cell division process. To determine the frequency of these rare cell division processes, we conducted QPI of expanded HSCs (Movie S4) over a wide field of view, measuring1000 μm. The imaging data captured 1243 cell division events within 36 hours, with 91.3% exhibiting normal division patterns, while 8.21% showed interruptions in cytokinesis, and 0.48% exhibited abnormal division patterns, such as a single cell dividing into three cells during one mitosis. (Fig. S1, D to G.)

**Fig. 1.**
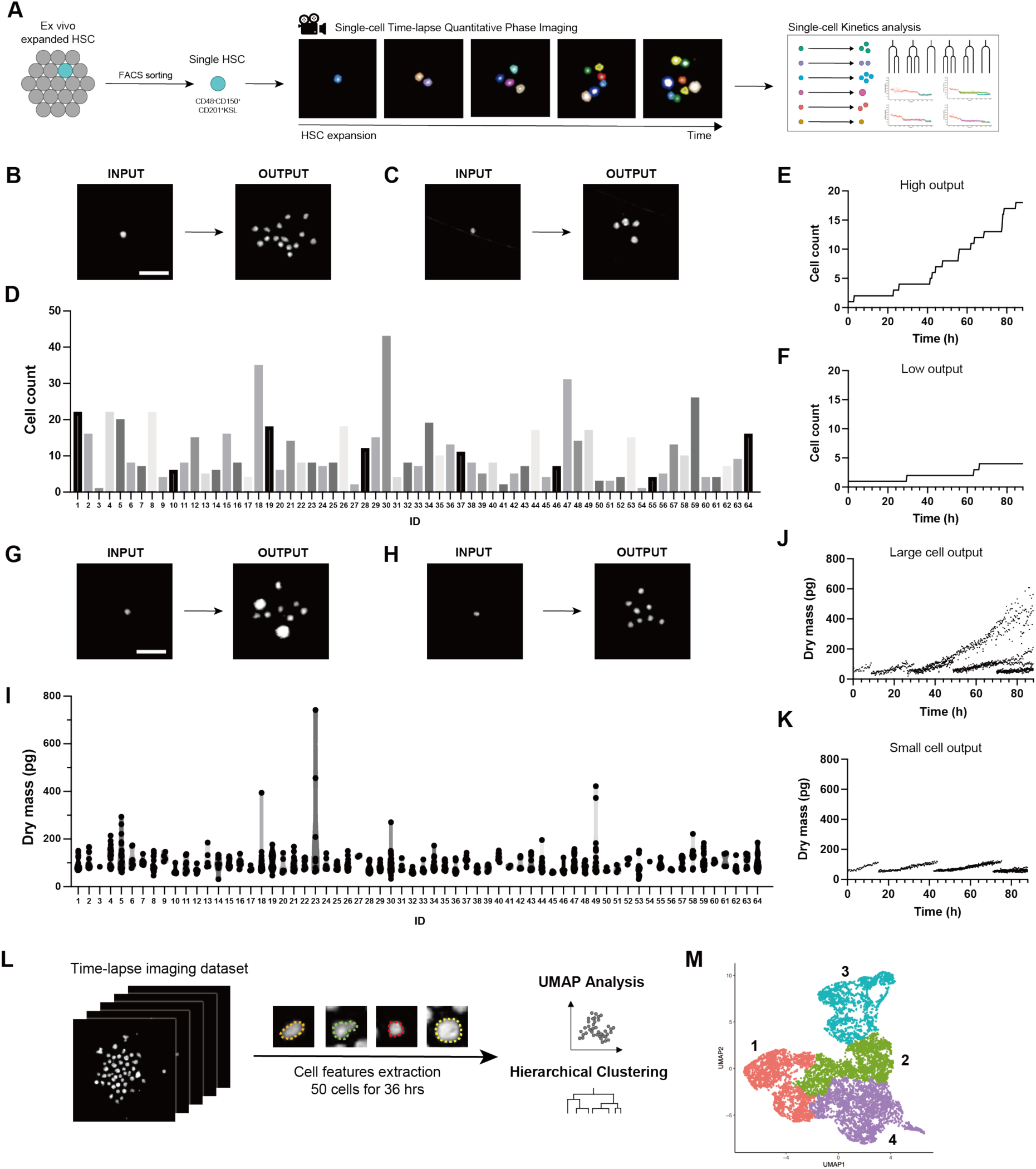
Quantitative phase imaging-based single-cell kinetic analysis of expanded HSCs. **(A**) Protocol for analyzing single HSC time-lapse quantitative phase imaging. A single CD48^-^ CD150^+^CD201^+^KSL ex vivo expanded HSC was sorted and monitored using time-lapse QPI for 96 hours. Cellular kinetics were analyzed quantitatively. (**B, C**) Representative images of HSCs that underwent rapid or slow divisions over 96 hours. Scale bar: 100 μm. (**D**) Number of cells produced by each HSC after 96 hours of expansion. Each ID represents a single HSC. (**E, F**) Representative proliferation patterns of HSCs that underwent rapid or slow divisions over 96 hours. (**G, H**) Representative images of HSCs that produce cells with high or low dry mass. (**I**) Dry mass of cells produced by each HSC after 96 hours of culture. Each ID represents a single HSC. (**J, K**) Representative dry mass kinetics of HSCs that produce either high- or low-mass cells. **(L)** Protocol for classifying cells based on kinetic features derived from QPI data. Fifty HSCs were tracked for 36 hours, and 11 parameters were extracted from 11,512 cell images for UMAP analysis and hierarchical clustering. **(M)** UMAP analysis of expanded HSCs based on kinetic features obtained by QPI, colored by hierarchical clustering.

Therefore, using QPI time-lapse imaging, individual HSCs were observed displaying unique behaviors even in pure phenotypic HSC fractions, highlighting the infinite diversity seen within the HSC population.

### Gene-independent cell classification through kinetic features of HSCs

Next, we evaluated the potential for cell classification based on kinetic features obtained from QPI data. We tracked 50 cells over 36 hours, and 11 parameters were extracted from a total of 11,512 cell images for UMAP analysis and clustering (Fig. 1L, Movie S5). This resulted in the identification of four distinct clusters, each with unique characteristics (Fig. 1M). Notably, Cluster 3 exhibited cells with low dry mass, high sphericity and low velocity, while Cluster 4 comprised cells with high dry mass (Fig. S2, B and C). This UMAP is derived from time-lapse imaging data and includes actual time information, enabling the tracking of cells on the UMAP. For example, Cell ID No.1 and No.2 remained within Cluster 3 for 36 hours without dividing, whereas Cell ID No.3 and No.4 underwent division and were found within Cluster 1,2 and 4. Cell ID No.5 remained in cluster 4 (Fig. S2D). By referencing the temporal data, we observed a temporal progression from the top of Cluster 3 through Cluster 1 and 2, and towards Cluster 4 (Fig. S2E). This suggests that cells at the top of Cluster 3 are the most immature HSCs.

### *Hlf* marks functional HSC during ex vivo expansion

Despite the identification of several genes expressed in fresh bone marrow HSCs, there is insufficient information on genes that are expressed specifically in ex vivo expanded murine HSCs. To capture the differentiation process of HSCs during ex vivo expansion, we first analyzed expanded HSCs on days 1, 3, 5 and 7 with flow cytometry, revealing that CD201^+^CD48^-^ cells gradually transitioned to CD201^+^CD48^+^ and CD201^-^CD48^+^ cells over time (Fig. 2, A and B, Fig. S3A). Reanalysis of bulkRNA-seq data for each fraction showed that *Hlf*, *Mecom* and *Fgd5* were highly expressed in the CD201^+^CD48^-^ fraction, with reduced expression in CD201^+^CD48^+^ and CD201^-^CD48^+^ cells (Fig. 2C, Fig. S3B) (*26*). Single-cell RNA sequencing of expanded HSCs revealed that *Hlf* was highly expressed in the HSC fraction and sharply decreased with differentiation (Fig. 2D, Fig. S3, C and D). Therefore, *Hlf* was determined to be an effective indicator of stemness levels during ex vivo HSC expansion.

**Fig. 2.**
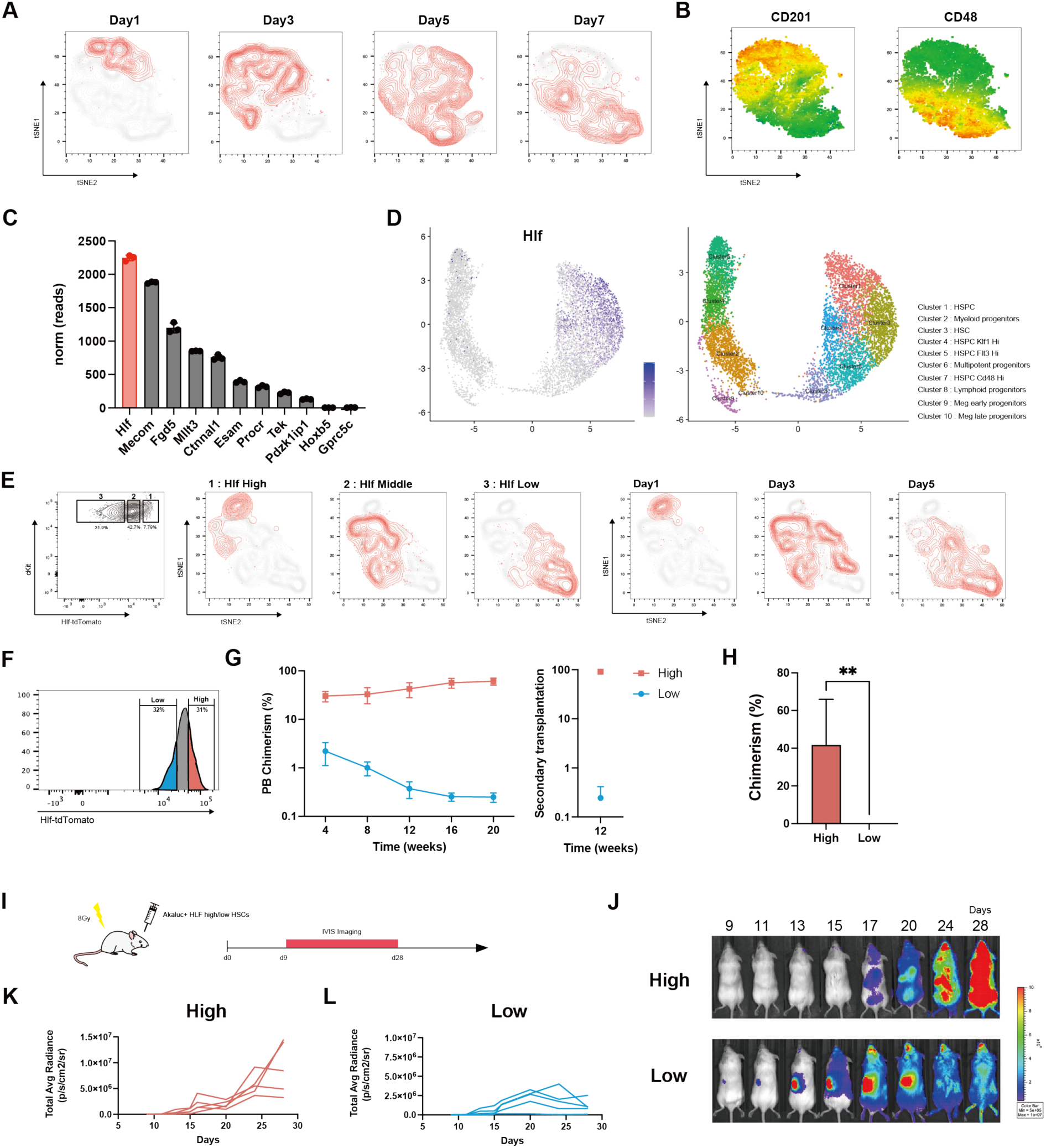
Hlf marks functional HSCs during ex vivo expansion. **(A)** tSNE analysis of ex vivo expanded HSCs based on FACS data using CD201, CD48, CD150, cKit, Sca1, and Lin markers. FACS data of HSCs after 1, 3, 5, and 7 days of expansion were combined for tSNE analysis. **(B)** tSNE representation of expanded HSCs, overlaid with CD201 and CD48 markers **(C)** RNA-seq expression profiles of selected HSC-associated genes in CD48^-^CD201^+^CD150^+^KSL cells (n = 3). Error bars represent standard deviation (SD). **(D)** UMAP plot of single-cell RNA-seq data from 7-day expanded HSCs with 10 annotated clusters and feature plots showing *Hlf* expression. **(E)** tSNE analysis of ex vivo expanded HSCs from Hlf-tdTomato mice. FACS data of HSCs after 1, 3, and 5 days of expansion were integrated for tSNE analysis. The leftmost plot shows the gating strategy for high (1), middle (2), and low (3) Hlf-tdTomato populations. The right plot highlights each population in red for different expansion days. **(F)** Gating strategy for Hlf-tdTomato high and low expanded HSCs. **(G)** Mean donor peripheral blood chimerism in primary recipients (n = 4, 5 mice per group) and secondary recipients (n = 3–5 mice per group). Error bars represent SD. **(H)** Mean donor bone marrow chimerism in primary recipients (n = 4, 5 mice per group). Error bars represent SD. Statistical significance was calculated using a t-test. **P<0.01 **(I)** Schematic illustrating the analysis of hematopoietic dynamics after Hlf-tdTomato high/low expanded HSC transplantation with AkaBLI. Luminescence signals were measured by IVIS from day 9 to day 28 post-transplantation. **(J)** Representative IVIS images from the Hlf-tdTomato high/low Akaluc+ HSC transplantation assay. **(K, L)** Dynamics of total average luminescence intensity after transplantation of Hlf-tdTomato high/low expanded HSCs.

Based on these results, we utilized Hlf-tdTomato reporter mice to quantitatively track the stemness level of HSCs (*30*). High expression of tdTomato was observed in bone marrow derived CD34^-^CD150^+^CD48^-^ KSL cells from Hlf-tdTomato reporter mice (Fig. S3E). We validated whether the fluorescence intensity of Hlf-tdTomato was an effective indicator of the differentiation process of HSCs in expansion culture by analyzing cells from day 1, 3 and 5 post-expansion with flow cytometry. As expected, a strong correlation between changes in Hlf-tdTomato expression and temporal differentiation in HSCs was confirmed (Fig. 2E, Fig. S3, F and G). To determine whether Hlf-tdTomato expression levels act as a functional indicator of HSCs, tdTomato high and low expanded HSCs were transplanted into irradiated recipients, and long-term bone marrow reconstitution abilities were evaluated. HSCs with high tdTomato demonstrate robust long-term marrow reconstitution capabilities. Conversely, chimerism decreased in those who received tdTomato-low HSCs (Fig. 2, F to H). Furthermore, the analysis of the spatiotemporal dynamics of early hematopoiesis post-transplantation revealed that tdTomato high HSCs gradually reconstituted systemic hematopoiesis, whereas tdTomato low HSCs induced rapid hematopoiesis between day 14 and 21 (Fig. 2, I to L).

Consequently, Hlf-tdTomato expression levels are a useful indicator of stemness in HSCs during ex vivo expansion and serve as a potent predictor of both long-term and short-term hematopoietic dynamics post-transplantation.

### Predicting *Hlf* expression levels from cellular kinetic features

Next, we analyzed whether kinetic features could predict stemness, using *Hlf* expression as a proxy indicator. We performed UMAP analysis using cellular kinetic features obtained from time-lapse imaging, excluding tdTomato fluorescence intensity, confirming multiple clusters resembling those observed in Figure 2 (Fig. 3, A and C, Fig. S4, A and B, Movie S6). Then the average fluorescence intensity of tdTomato was plotted onto the same plot, which revealed significant differences in tdTomato expression levels among clusters (Fig. 3, B and D). In particular, we observed the highest tdTomato expression in Cluster 1 that gradually decreased from Cluster 2 through Cluster 5 (Fig. 3, B and D). This result was validated using other data sets (Fig. S4, C and D). Cells with high tdTomato expression exhibited higher sphericity, lower dry mass, and lower velocity than those with low tdTomato expression (Fig. 3E). These findings suggest that Hlf-tdTomato expression levels can be predicted from kinetic features of HSCs.

**Fig. 3.**
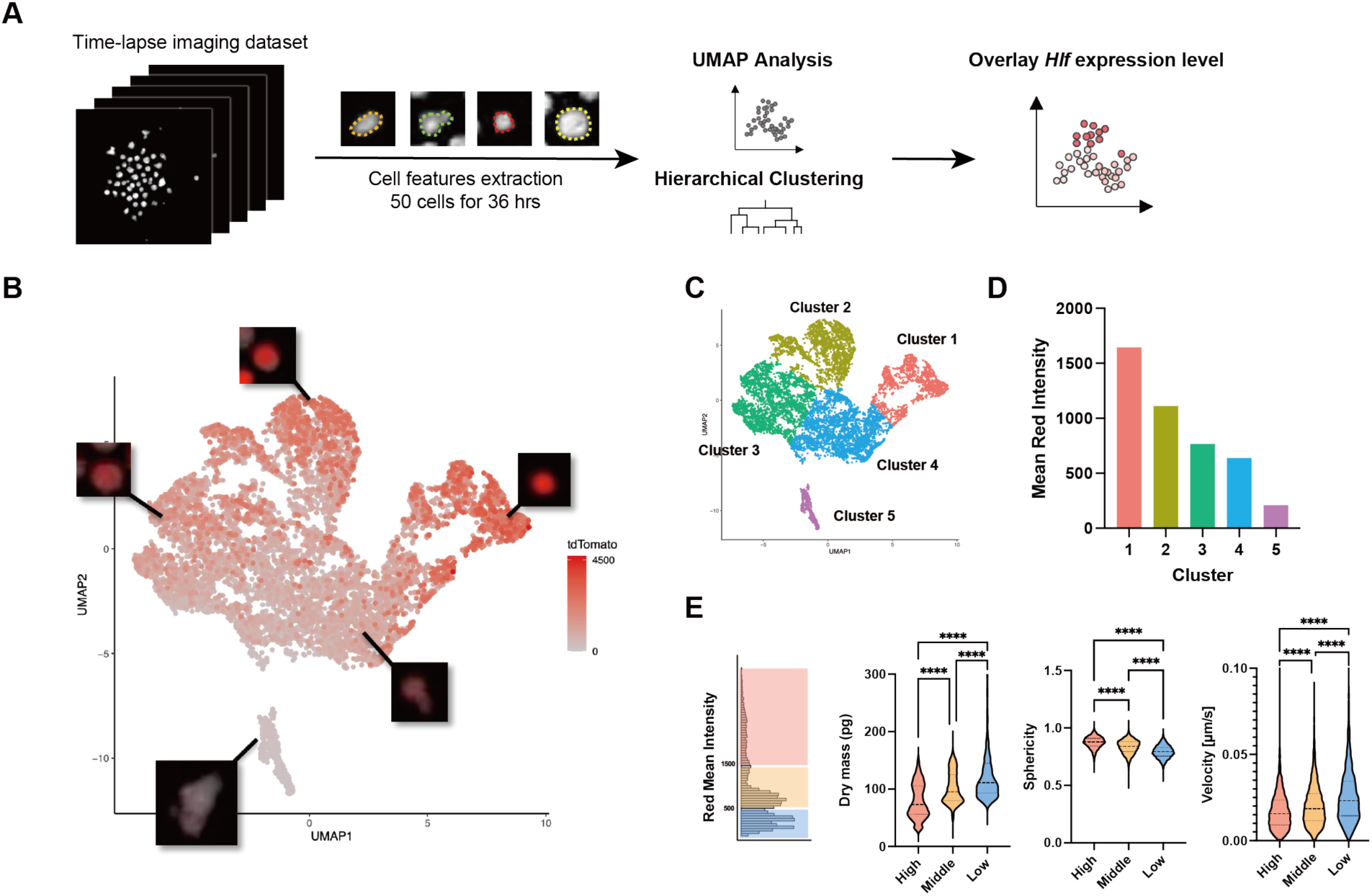
Predicting Hlf expression levels from cellular kinetic features. **(A)** Protocol for cell classification based on kinetic features from QPI data. Fifty HSCs were tracked for 36 hours, and 11 parameters were extracted from 11,876 cell images for UMAP analysis and hierarchical clustering, followed by overlay of Hlf-tdTomato expression level based on fluorescence imaging. **(B)** UMAP plot of expanded HSCs based on kinetic features, overlaid with Hlf-tdTomato expression level. Representative images of clusters are shown on the plot. **(C)** UMAP plot colored by hierarchical clustering. **(D)** Mean red intensity of each cluster. **(E)** Violin plot comparing dry mass, sphericity, and velocity among Hlf-tdTomato high, middle, and low expanded HSCs. Error bars represent SD. Statistical significance was calculated using one-way ANOVA with Tukey’s post-test. ****P<0.0001

### Diversity of HSCs based on single cell kinetic analysis with *Hlf* dynamics

To analyze individual cell kinetics more precisely, we performed index sorting on the CD201^+^CD48^-^ CD150^+^KSL fraction of expanded Hlf-tdTomato^+^ HSCs, followed by ex vivo expansion and 96-hour QPI (Fig. 4A). We observed diverse cellular kinetics between every single cell. By tracking changes in Hlf-tdTomato expression levels, we confirmed that diversity exists in fluorescence intensity dynamics among cells. After 96 hours of expansion, 18.8% of HSCs produced high Hlf-tdTomato cells (>5000), whereas 14.1% of HSCs produced only low Hlf-tdTomato cells (<1000) (Fig. 4B). The index sorting data revealed a positive correlation between the number of output cells and CD150 expression levels (P<0.0001, R squared = 0.40) (Fig. 4L) and a weak negative correlation with Hlf-tdTomato levels (P=0.0072, R squared = 0.11) (Fig. 4M). The average intensity of Hlf-tdTomato in output cells was positively correlated with Hlf-tdTomato (P<0.0001, R squared = 0.53) (Fig. 4N) and other markers in input cells, such as CD201, Sca1 and cKit (Fig. S5A), while negatively correlated with the number of output cells (P<0.0001, R squared = 0.38) (Fig. 4O). Cells that divided quickly showed a decrease in Hlf-tdTomato expression over time, while those that divided slowly maintained Hlf-tdTomato levels (Fig. 4, C and D, F and G, I and J, Fig. S5, B and C, Movie S7). In fact, Hlf-tdTomato high HSCs predominantly produced HSC-like cells, whereas Hlf-tdTomato low HSCs produced more differentiated downstream cells (Fig. S5, D to G). Some cells maintained high Hlf-tdTomato expression and divided, but eventually led to early cell death (Fig. 4, E, H and K, Movie S8). These cells showed lower levels of CD150 expression. To investigate the in vivo dynamics of these cells, CD150^high^ Hlf-tdTomato^high^ CD201^+^CD48^-^KSL cells and CD150^dim^ Hlf-tdTomato^high^ CD201^+^CD48^-^KSL cells were transplanted into irradiated mice. As a result, CD150^high^ cells showed long-term reconstitution capabilities, whereas CD150^dim^ cells exhibited short-term reconstitution abilities, and the chimerism gradually decreased over time (Fig. S5, H to K). Thus, Hlf-tdTomato^high^ CD150^dim^ cells maintained Hlf-tdTomato expression ex vivo but exhibited short-lived characteristics, indicating short-term HSCs.

**Fig. 4.**
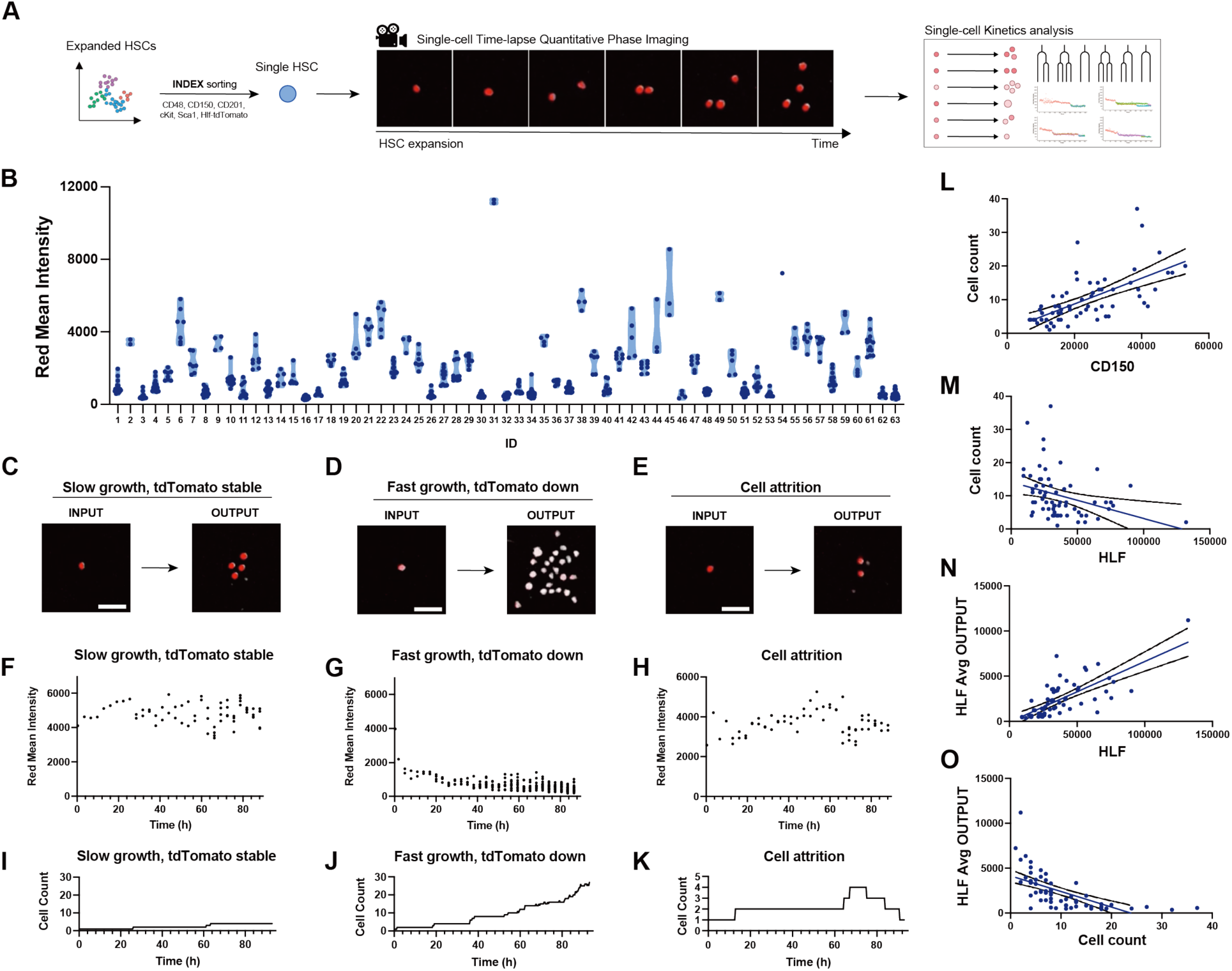
Single HSC kinetics analysis with Hlf-tdTomato dynamics. **(A)** Protocol for single HSC time-lapse QPI. A single CD48^-^CD150^+^CD201^+^KSL ex vivo expanded HSC was sorted and monitored using time-lapse QPI for 96 hours. Cellular kinetics were quantitatively analyzed. **(B)** Red mean fluorescence intensity of each cell produced after 96 hours. **(C-E)** Representative images of single sorted HSCs (input) and their produced cells (output) over 96 hours. Scale bar: 100 μm. Hlf-tdTomato expression shown as red overlay. **(F-K)** Representative kinetics of HSCs that maintained high Hlf-tdTomato levels and divided slowly (F, I), decreased Hlf-tdTomato levels and proliferated rapidly (G, J), or maintained Hlf-tdTomato levels but did not proliferate (H, K). **(L)** Correlation between the CD150 level of sorted single HSCs and the cell count after 96 hours. Simple linear regression (blue line) and 95% CI (black lines). **(M)** Correlation between the Hlf-tdTomato level of sorted single HSCs and the cell count after 96 hours. Simple linear regression (blue line) and 95% CI (black lines). **(N)** Correlation between the Hlf-tdTomato level of sorted single HSCs and their produced cells after 96 hours. Simple linear regression (blue line) and 95% CI (black lines). **(O)** Correlation between the cell count and Hlf-tdTomato level of produced cells after 96 hours. Simple linear regression (blue line) and 95% CI (black lines).

Despite the general trends observed, individual cell kinetics showed significant variations. Some cells maintained Hlf-tdTomato expression while producing many cells, indicating high self-renewal capabilities as HSCs (Fig. S5K, Movie S9). However, these cells could not be identified using index sorting data. The size of output cells also varied, showing a weak negative correlation with cKit, though this correlation coefficient was low and not statistically significant, making predictions using index sorting data challenging (Fig. S5A). Additionally, no correlation was found between Division Gap and index sorting data (Fig. S5A). Thus, through the integration of QPI with temporal changes in Hlf-tdTomato expression, we were able to gain deeper insight into the complexity and diversity of HSCs beyond what can be captured by FACS analysis alone.

### Machine learning prediction of *Hlf* expression levels from live-cell behavioral kinetics

We have demonstrated the feasibility of predicting Hlf-tdTomato expression levels in HSCs using UMAP analysis with multiple cellular kinetic features extracted from QPI data. However, there are only a limited number of biological features used in the analysis. Therefore, we developed a system that can more accurately predict the *Hlf* expression levels of each cell by training a deep neural network with QPI datasets. We tracked individual target cells and extracted QPI video data specific to the target cells, generating a dataset that matched the video data of each cell from Frame 1 to Frame n with the corresponding Hlf-tdTomato intensity at the final frame (Frame n), thereby excluding the influence of surrounding cells from the videos. Machine learning was conducted using 3D Residual Neural Network (ResNet) on this training dataset, and the model was validated using a separate dataset (Fig. 5A). As a result, when predictions were made utilizing a sequence of 20 consecutive frames, the expression level of Hlf-tdTomato was predicted with remarkably high accuracy (correlation coefficient: 0.73, error: 0.02) (Fig. 5, C to E). Notably, by increasing the number of frames employed for learning from 5 to 20, the prediction accuracy significantly improved, more than doubling (Fig. 5E).

**Fig. 5.**
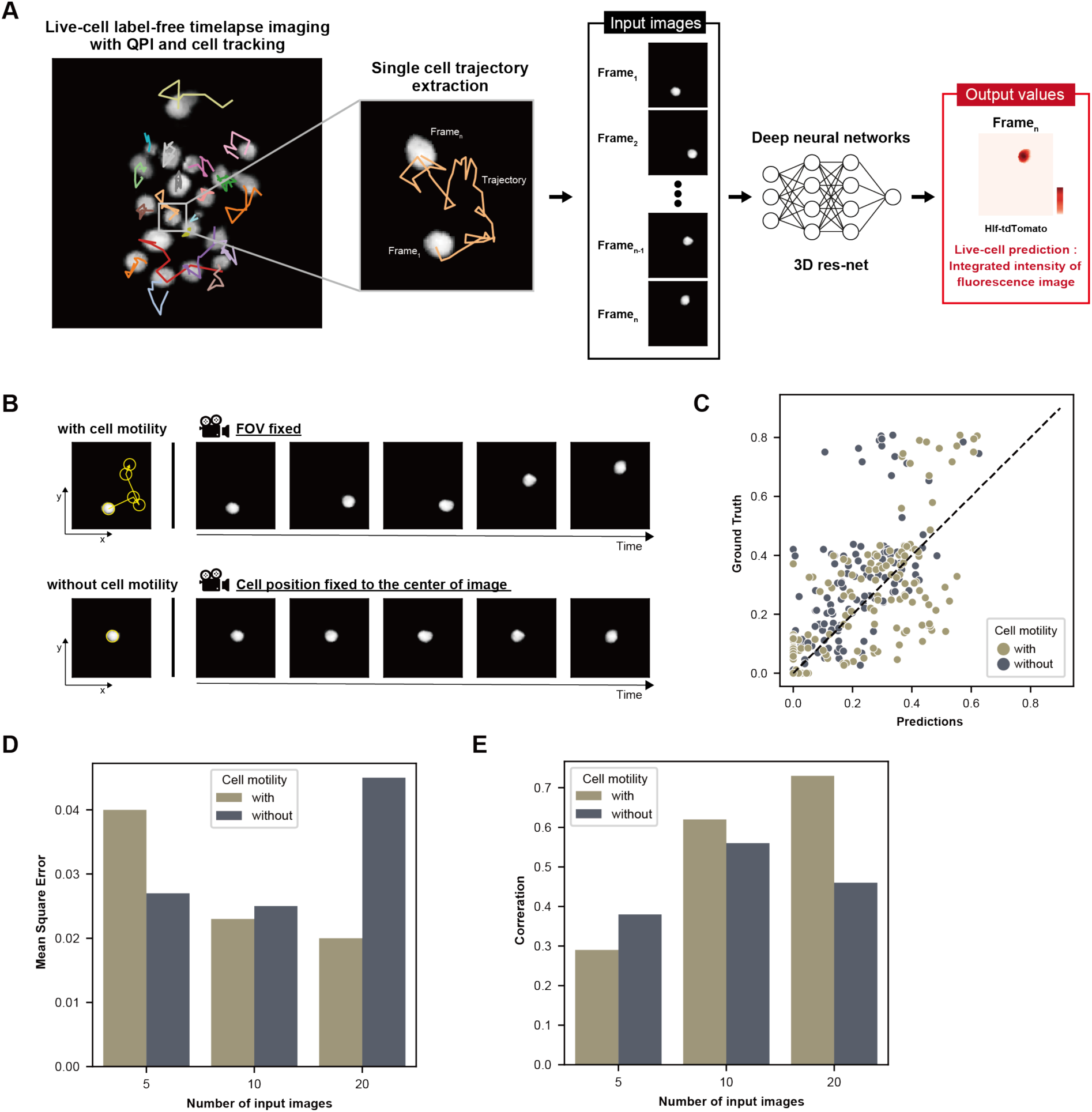
QPI-driven machine learning prediction of *Hlf* expression levels. **(A)** Protocol for QPI-driven machine learning to predict the Hlf-tdTomato level of live HSCs. Time-lapse imaging was conducted every 4 minutes and 21 seconds. Individual cells were tracked, and their trajectories were extracted to generate a dataset matching video data of each cell from frame 1 to frame n with the corresponding Hlf-tdTomato intensity in the final frame (frame n). 3D ResNet was used on this training dataset. **(B)** Illustration of representative datasets with fixed field-of-view images (above) and images centered on cell position (below). **(C)** QPI-driven machine learning with and without cell motility, comparing predicted Hlf-tdTomato expression levels (x-axis) and measured Hlf-tdTomato levels (y-axis). **(D)** Mean squared error of Hlf-tdTomato predictions with and without cell motility using 5, 10, and 20 input images. **(E)** Correlation coefficient of Hlf-tdTomato predictions with and without cell motility using 5, 10, and 20 input images.

We also investigated the impact of cellular kinetics on prediction accuracy. Previous analysis had shown that less migratory HSCs tended to express high levels of *Hlf* (Fig. 3). To assess the impact of HSC motility on prediction accuracy, we removed the X, Y positional information from the tracked cells, fixing the position of cells in the center of the video, and made prediction of Hlf-tdTomato expression levels (Fig. 5B, Movie S10). Consequently, when positional information was excluded, the prediction accuracy of Hlf-tdTomato expression levels significantly decreased and increasing the number of frames did not improve accuracy (Fig. 5, C to E). This indicates that the motility of HSCs during ex vivo expansion is an important indicator of stem cell properties.

In conclusion, integrating HSC ex vivo expansion technologies with QPI to continuously monitor HSC characteristics in a temporally-dependent manner reveals new HSC diversity previously undetectable by snapshot analysis. Furthermore, the use of a predictive model leveraging multidimensional analysis based on cellular kinetic features and a deep neural network has demonstrated that the introduction of temporal information significantly enhances the accuracy of predicting stem cell properties from QPI data. Together, these achievements highlight the importance of the temporal dimension in uncovering the dynamic behavior and functional heterogeneity of HSCs.

### Concluding Remarks

In this study, we developed a system that continuously tracks the kinetic features and dynamics of HSCs by integrating our established HSC expansion culture system with QPI technology, allowing for high-precision, label-free prediction of HSC function. By increasing the number of video frames and considering the temporal mobility of cells, the system achieves a significant improvement in accuracy, highlighting the importance of evaluating cellular kinetic features over time. This technology revolutionizes the approach to HSC identification and establishes a next-generation label-free functional prediction platforms for expanded HSCs.

Through multidimensional analysis using extracted feature metrics, we have successfully classified cells into multiple categories, independent of genetic information, and have charted their continuous cell differentiation trajectories on UMAP. This strategy enables the analysis of authentic temporal shifts in the same cell, in contrast to estimated temporal data, such as pseudo time analysis obtainable through scRNAseq. Indeed, incorporating temporal information revealed diversities in HSCs that remains obscured in traditional snapshot analysis. We have recently developed a novel culture system for human HSCs and anticipate that the same prediction system applied to murine models can be adapted to human cells (*27*).

The system developed in this study enables the non-invasive prediction of HSC stemness at a single-cell level. To directly validate the system, the ideal methodology would involve harvesting and transplanting HSCs at the single-cell level. Although currently unfeasible due to technological barriers, it is anticipated that future advancements in technology will enable proof of this concept.

## Acknowledgments

We thank the University of Tokyo Institute of Medical Science (IMSUT) FACS core laboratory for technical assistance. We thank Dr. Masafumi Muratani and Tsukuba i-Laboratory LLP for sequencing services. We further thank Dr. Martin Humphry of Phasefocus for providing expert knowledge regarding quantitative phase imaging.

## Author contributions

Conceptualization: TY, SY

Methodology: TY, SY

Investigation: TY, YI, HJB, TK

Visualization: TY, YI

Funding acquisition: SY

Project administration: SY

Supervision: TY, TS, SO, SY

Writing – original draft: TY

Writing – review & editing: TY, YI, HJB, ASF, TK, TY, TS, SO, SY

## Data and materials availability

All data are available in the main text or the supplementary materials. All data have been uploaded to a Mendeley Data repository and will be published upon acceptance of the manuscript (DOI: 10.17632/hzvpmrf5g3.1). Further information and requests for resources and reagents should be directed to and will be fulfilled by TY or SY.

## Code Availability

Code for all our representation learning models is available at https://github.com/Iwamo-yu/QPI-HSCs.

## Materials and Methods

### Mice

C57BL/6NCrSlc (Ly 5.2, CD45.2) mice were purchased from SLC Inc., Japan. C57BL/6-Ly5.1 (Ly5.1, CD45.1) mice were obtained from Sankyo Labo Service Corporation, Inc., Japan. C57B/6 albino mice were purchased from The Jackson Laboratory Inc., Japan. All mice were obtained at the age of 8-10 weeks and maintained in a specific-pathogen-free environment with free access to food and water. Hlf-tdTomato mice were kindly provided by Tomomasa Yokomizo (Department of Microscopic and Developmental Anatomy, Tokyo Women’s Medical University, Tokyo, Japan.). All animal studies were conducted in accordance with institutional protocols and were approved by the Animal Care and Use Committee of the Institute of Medical Science at the University of Tokyo and the Laboratory Animal Resource Center at the University of Tsukuba.

### Murine HSC isolation

Male 8–10-week-old C57BL/6-CD45.1 mice were humanely sacrificed under isoflurane anesthesia. Pelvic, femur, and tibia bones were excised and crushed, and the resulting cell solution was filtered and the whole bone marrow cells were enumerated. Positive selection of cKit+ cells was accomplished using anti-APC magnetic-activated cell sorting (MACS, Miltenyi Biotec) antibodies after staining the cells with cKit-APC antibody (Thermo Fisher Scientific) for 30 minutes. Enriched cKit+ cells were then incubated with an anti-Lineage antibody cocktail (including biotinylated Gr1[LY-6G/LY-6C], CD11b, CD4, CD8a, CD45R[B220], IL7-R, TER119 (all from Thermo Fisher Scientific)) for an additional 30 minutes. This was followed by a 90-minute incubation with CD34-FITC (Thermo Fisher Scientific), Sca1-PE (Thermo Fisher Scientific), cKit-APC, streptavidin-APC/eFluor (Thermo Fisher Scientific), CD150-PE/Cy7 (BioLegend) and CD48-PB (BioLegend) antibodies. Propidium iodide (PI) was employed to exclude dead cells. CD34^-^CD150^+^ CD48^-^ cKit^+^Sca1^+^Lin^-^ (CD34^-^CD150^+^ CD48^-^KSL) cells were sorted via fluorescence-activated cell sorting (FACS) on Aria III cell sorter (BD) using a 100 um nozzle with appropriate filters and settings. For Hlf-tdTomato reporter mice, cKit+ cells were incubated with an anti-Lineage antibody cocktail, followed by incubation with CD34-FITC, Sca1-BV605 (BioLegend), cKit-APC, streptavidin-APC/eFluor, CD150-PE/Cy7 and CD48-PB (BioLegend) antibodies. CD34^-^CD150^+^CD48^-^cKit^+^Sca1^+^Lin^-^tdTomato^+^ (CD34^-^CD150^+^CD48^-^ Hlf^+^KSL) cells were sorted via FACS on Aria III cell sorter.

### Murine HSC culture

Murine hematopoietic stem cells (HSCs) were cultured in Ham’s F12 medium (Wako), enriched with 10 mM HEPES (Thermo Fisher Scientific), recombinant cytokines murine TPO (100 ng/mL, Peprotech), and SCF (10 ng/mL, Peprotech), Soluplus (BASF), as well as insulin-transferrin-selenium (ITS, Thermo Fisher Scientific, 1:100 dilution) and 1% Penicillin-Streptomycin-L-Glutamine (PSG, Wako). Murine HSCs were cultured on untreated U-bottom or flat-bottom 96-well plates (TPP, for cultures starting with 100 cells). Cells were maintained in an incubator (Panasonic) at 37℃ with a constant CO_2_ fraction of 5%, and medium changes were carried out every 2-3 days.

### Quantitative phase imaging and analytical tools

Quantitative phase imaging was conducted with Livecyte (Phasefocus) on U-bottom/flat-bottom 96 well plate. Cells were maintained at 37℃ with a constant CO_2_ fraction of 5%. Phase and fluorescence images were acquired in parallel for each well. In single-cell imaging, imaging was started 8 hours after sorting. In bulk-cell imaging, measurements were started 15 hours after sorting. Phasefocus’ Cell Analysis Toolbox software was used for cell segmentation, cell tracking, and data exportation in phase images. Segmentation thresholds were optimized using various image processing techniques, such as the rolling ball algorithm for background noise removal, image smoothing for detecting cell edges, and local pixel maxima detection to identify seed points for final consolidation. The feature tables output by the Phasefocus software for each imaged well were analyzed using R. For cell division pattern analysis, Trackmate ImageJ plugin was used to identify cell divisions and classify the division patterns. For the UMAP analysis based on cellular kinetic characteristics, parameters such as Volume, Thickness, Radius, Area, Sphericity, Length, Width, Length-width ratio, Dry mass, Velocity, and Perimeter were extracted from the feature table

### Peripheral blood analysis

Peripheral blood was collected using a heparin tube. For chimerism and lineage analysis, erythrocytes were lysed in NH_4_Cl solution. The resulting lysed blood cells were stained with Gr1-PB (BioLegend), CD11b-PB (BioLegend), CD4-APC (Thermo Fisher Scientific), CD8a-APC (Thermo Fisher Scientific), CD45R[B220]-APC/eFluor 780 (Thermo Fisher Scientific), CD45.1-PE/Cy7 (Tombo Biosciences) and CD45.2-BV421 (Thermo Fisher Scientific) for C57BL/6 mice samples. The stained cells were then resuspended in 200 μl PBS/PI prior to recording events on a FACSVerse (BD) analyzer using the appropriate filters and settings.

### Cell counting and sample preparation for flow cytometry

Cell counting was carried out utilizing an automated cell counter (Countess II cytometer, Thermo Fisher Scientific). For hematopoietic stem cells in post-transplant bone marrow, cells were incubated with an anti-Lineage antibody cocktail consisting of biotinylated Gr1[LY-6G/LY-6C], CD11b, CD4, CD8a, CD45R[B220], IL7-R, and TER119. This was followed by staining with streptavidin-APC/eFluor, cKit-APC, Sca1-BV605 (BioLegend), CD150-PE/Cy7, and CD34-FITC. For sorting of ex vivo expanded HSCs after culture, cultured bulk cells were stained with an anti-Lineage antibody cocktail consisting of biotinylated Gr1[LY-6G/LY-6C], CD11b, CD4, CD8a, CD45R[B220], IL7-R, and TER119, followed by staining with streptavidin-BV421 (BioLegend), cKit-APC, Sca1-APC/Cy7 (BioLegend), CD150-PE/Cy7, and CD201-PE (Thermo Fisher Scientific) for cultured HSCs, and streptavidin-APC/eFlour, cKit-BV421, Sca1-APC/Cy7, CD150-PE/Cy7, CD201-APC (Thermo Fisher Scientific) and CD48-FITC for cultured Hlf-tdTomato HSCs. The stained cells were then resuspended in 200 μl PBS/PI and analyzed using FACS Verse analyzer and FACS AriaIII cell sorter.

### Akaluc vector transduction

Cultured cells underwent transduction with a VSV-G pseudotyped lentiviral vector carrying an mNeonGreen-P2A-Akaluc transgene under the regulation of the human ubiquitin C (UbC) promoter at a multiplicity of infection (MOI) of 300. A medium change was performed one day after transduction, and further medium changes were performed every 2-3 days.

### In vivo luminescence imaging

The mice were anesthetized with isoflurane and injected with 50 µL of TokeOni (15 mM, Kurogane Kasei Co., Ltd.) intraperitoneally. They were then placed in an IVIS in vivo imaging system (PerkinElmer). Images were taken after 5 minutes using appropriate binning and exposure settings. The luminescence signal intensities were analyzed with Living Image Software. To measure the luminescence signal intensity of the whole body, the ROI position for each mouse was kept consistent, and the temporal changes in average luminescence intensity at the same position were analyzed.

### Transplantation assay

Cell transplantation via intravenous injection was conducted in irradiated (8.0 Gy) C57BL/6-CD45.1 recipient mice, along with 5×10^5^ C57BL/6-CD45.1/CD45.2 whole bone marrow competitor cells, using cultured cells at indicated cell doses. Secondary bone marrow transplantations were performed by extracting WBM cells from the primary recipient and transplanting 1×10^6^ cells into lethally irradiated secondary recipients.

### Training Dataset Preparation Through Segmentation and Tracking

Our methodology started by capturing single-cell tracking videos with a quantitative phase microscope, which we then used as the input for our prediction model. Initially, we segmented each video frame using Voronoi Otsu labeling to identify each cell in every frame. Following this segmentation, we generated a tracking video for each cell. This step involved aligning the centroid of each cell across successive frames using the segmentation labels. We utilized Phase Focus’s live cell analysis software to create two types of videos: one that recorded cell movements within a static field of view (unchanged from the first frame) alongside their morphological changes and another that focused solely on morphological changes, cropping each frame to keep the cells centered. We manually verified every cell within the dataset to ensure data integrity.

We determined the prediction model’s output by measuring the cells’ cumulative fluorescence intensities. This process used the segmentation labels from the phase images as masks to measure the cumulative fluorescence intensity of cells in the video’s last frame, which we then used as the target for regression analysis.

### Prediction Model Training

We employed the 3D ResNet18 model for training, coding the regression task with the Python-based library Pytorch Lightning to predict the fluorescence intensity of cells from video data. Our dataset consists of videos from three independent experiments for training, with videos from a fourth experiment set aside for validation. We conducted cross-validation within the training dataset, dividing it into three subsets using the Stratified K-Folds cross-validation method to ensure representativeness in each fold. We chose the mean square error as the loss function to optimize this model due to its effectiveness in regression tasks.

### Single-cell RNAseq analysis

Murine bone marrow derived CD34^-^KSL cells were cultured for 7 days and collected. Libraries were generated using Single Cell 3’ Reagent Kit v.3.1 (10x Genomics), and sequenced on Illumina Hiseq X. Sequence data were mapped to reference genome (mm10) using CellRanger. Subsequent analysis was performed using Seurat R package. We filtered out cells that had nFeature_RNA over 5,000 or less than 200, and percentage of mitochondrial counts higher than 7.5%. Data was normalized and scaled using NormalizeData and ScaleData function. To reduce dimensions, Principal component analysis (PCA) and UMAP were performed using RunPCA and RunUMAP function. To identify clusters, we performed FindClusters function (resolution = 0.4).

### Bulk RNAseq analysis

In our bulk RNA-seq analysis, we performed a reanalysis of data we had obtained previously (26). Briefly, the target cells were sorted into 1.5 mL tubes and lysed in 600 µL Trizol LS reagent (Thermo Fisher Scientific). Subsequently, RNA purification, library preparation, and next-generation sequencing were performed by Tsukuba i-Laboratory, LLC. Libraries were generated using the SMARTer cDNA synthesis kit (Takara) and the high-output kit v2 (Illumina), followed by sequencing on a NextSeq 500 sequencer (Illumina) with 2×36 paired-end reads. DESeq2 package in R was utilized for data normalization and comparative analysis.

### Quantification and statistical analysis

Information concerning the statistical tests applied, the number of subjects and groups are mentioned in the figure legends. Student’s t-tests, one- and two-way analysis of variance (ANOVA) were executed using Prism (version 9.5, Graphpad). Standard deviations are represented by error bars. For statistical analysis related to RNAseq, R version 4 (R Core Team, 2020) along with the fitting package were utilized. Illustrations and diagrams were produced utilizing BioRender (https://www.biorender.com/).

## SUPPLEMENTAL FIGURE TITLES AND LEGENDS

**Fig. S1.**
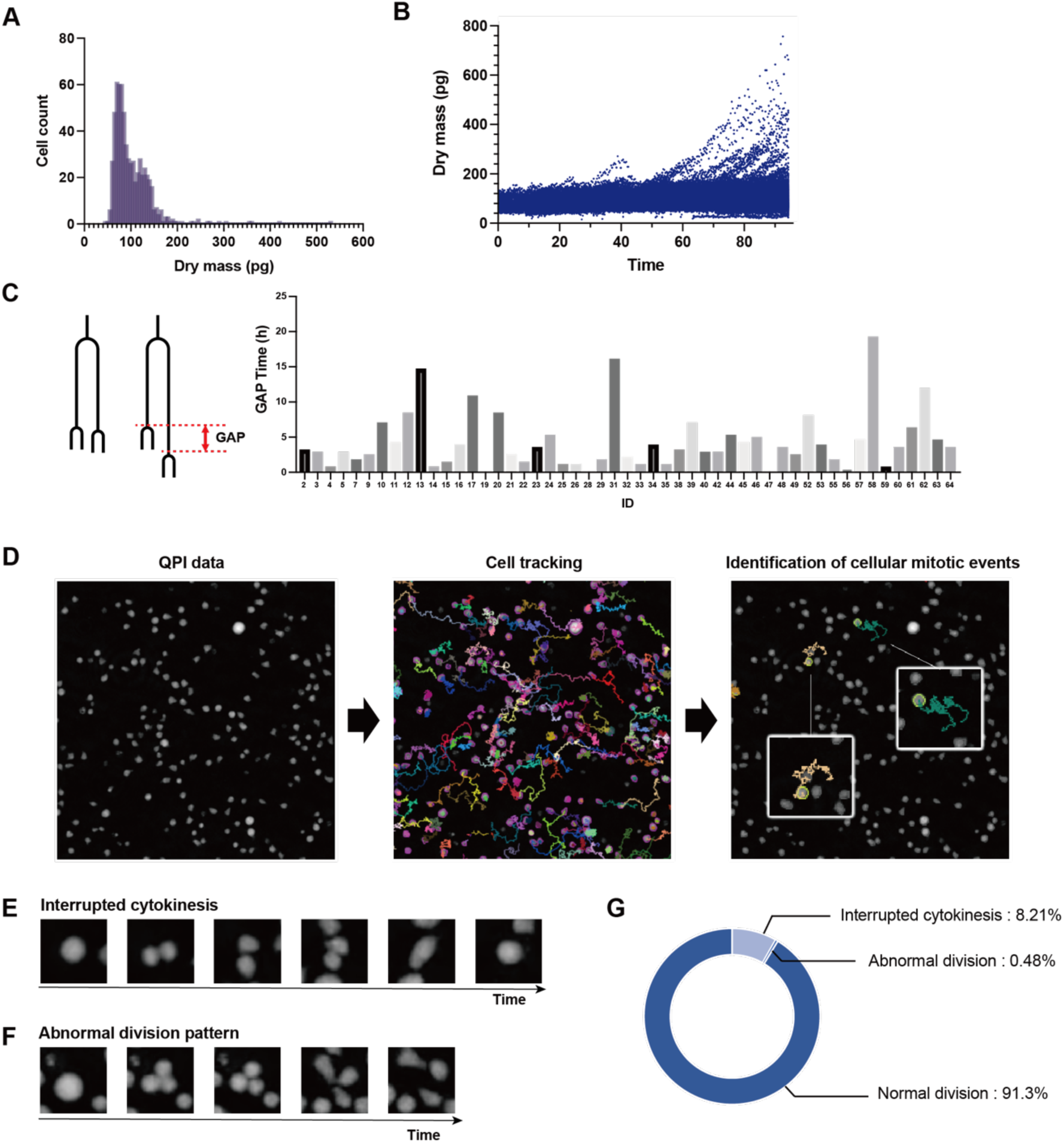
Cellular kinetic analysis based on quantitative phase imaging. **(A)** Histogram of dry mass of expanded HSCs after 96 hours of single-cell culture. All data were integrated. **(B)** Dynamics of HSC dry mass during single-cell expansion. All data were integrated. **(C)** Division gap for each HSC during single-cell expansion. Each ID refers to an individual HSC. **(D)** Protocol for extracting cell divisions. Fifty CD34^-^CD150^+^KSL HSCs were expanded over 7 days, and QPI was conducted for 36 hours. All cells were tracked, and cell division events were identified. **(E, F)** Representative rare division patterns: interrupted cytokinesis (E) and abnormal division pattern (F). **(G)** Frequency of each cell division type including abnormal divisions.

**Fig. S2.**
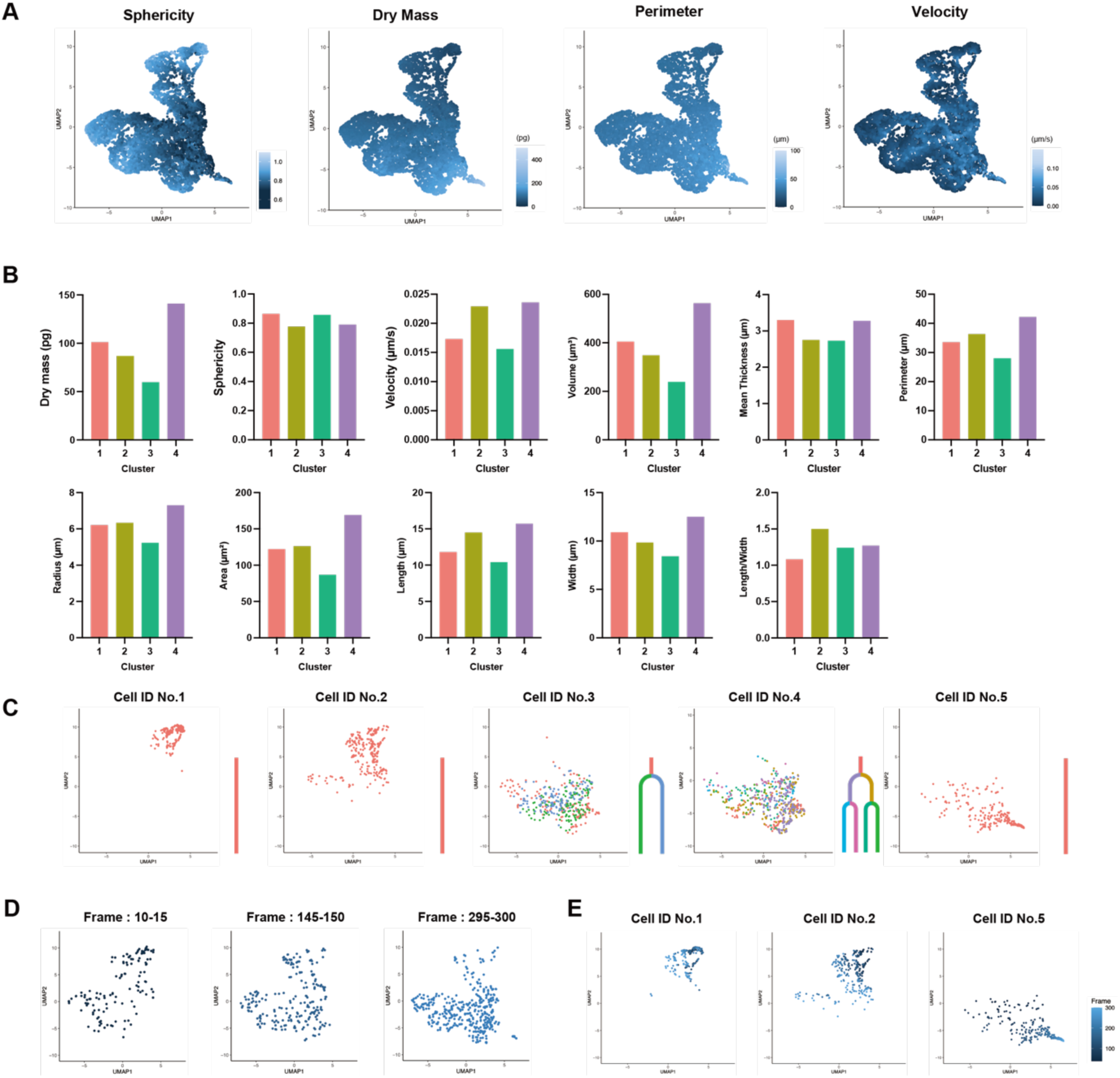
Single-cell kinetic feature analysis of expanded HSCs and cell classification. **(A)** UMAP representation overlaying sphericity, dry mass, perimeter, and velocity. **(B)** Average values of each parameter per cluster used in the analysis. **(C)** Representative cell tracking on the UMAP plot. Each color represents a distinct division. **(D)** UMAP representation of expanded HSCs during frames 10-15, 145-150, and 295-300. **(E)** UMAP representation of tracked HSCs overlaid with frame data.

**Fig. S3.**
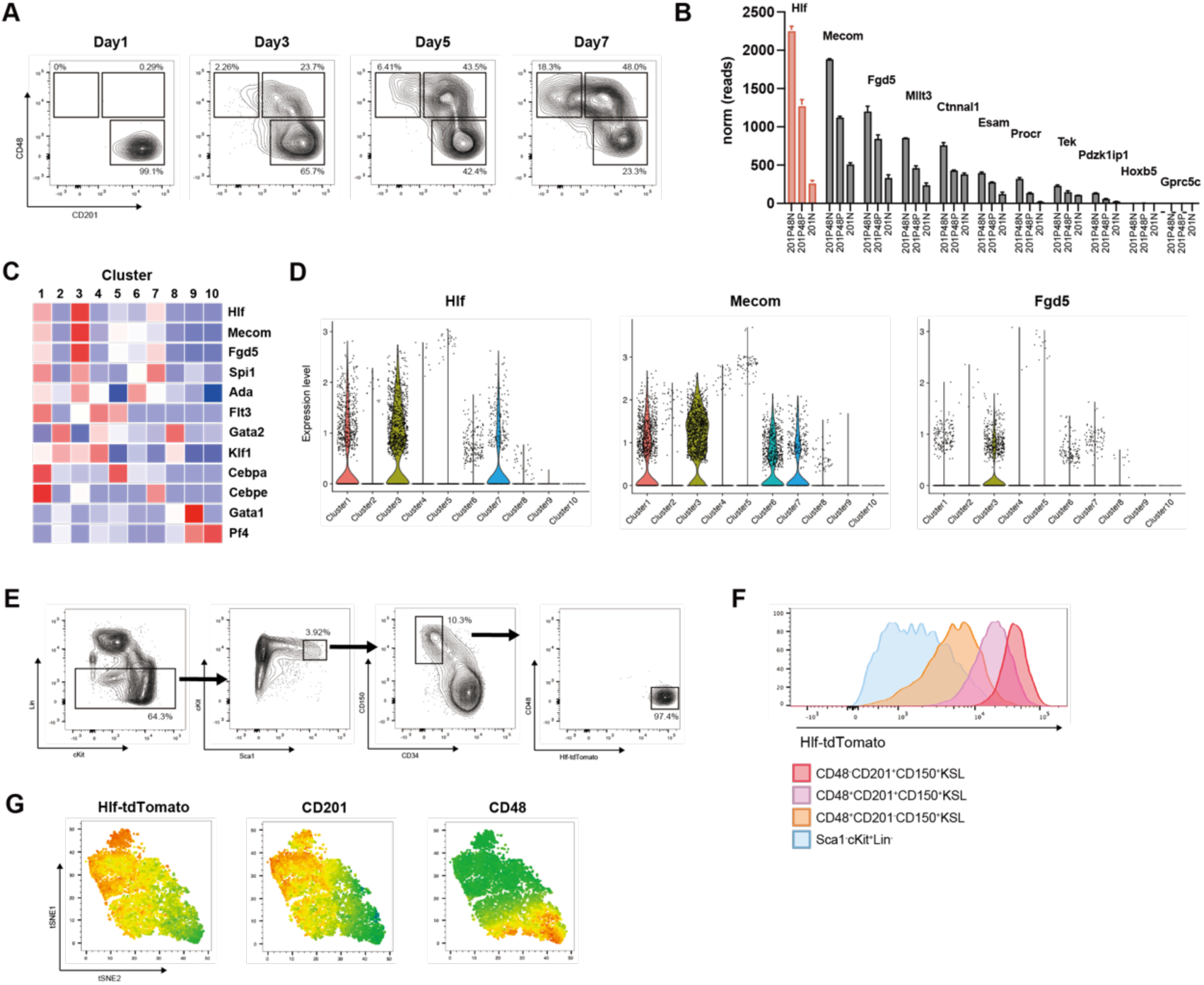
Hlf expression levels as an indicator of stemness in HSCs during ex vivo expansion. **(A)** FACS plots of expanded HSCs after 1, 3, 5, and 7 days. **(B)** RNA-seq expression profiles of select HSC-associated genes (n=3). Error bars represent standard deviation. **(C)** Heatmap analysis of single-cell RNA-seq data from 7-day expanded HSCs. **(D)** Violin plot of selected HSC-associated genes. **(E)** Gating strategy for Hlf-tdTomato^+^ fresh HSCs. **(F)** Comparison of Hlf-tdTomato levels among HSPC fractions. **(G)** tSNE plot of expanded Hlf-tdTomato HSCs colored by Hlf-tdTomato, CD201, and CD48.

**Fig. S4.**
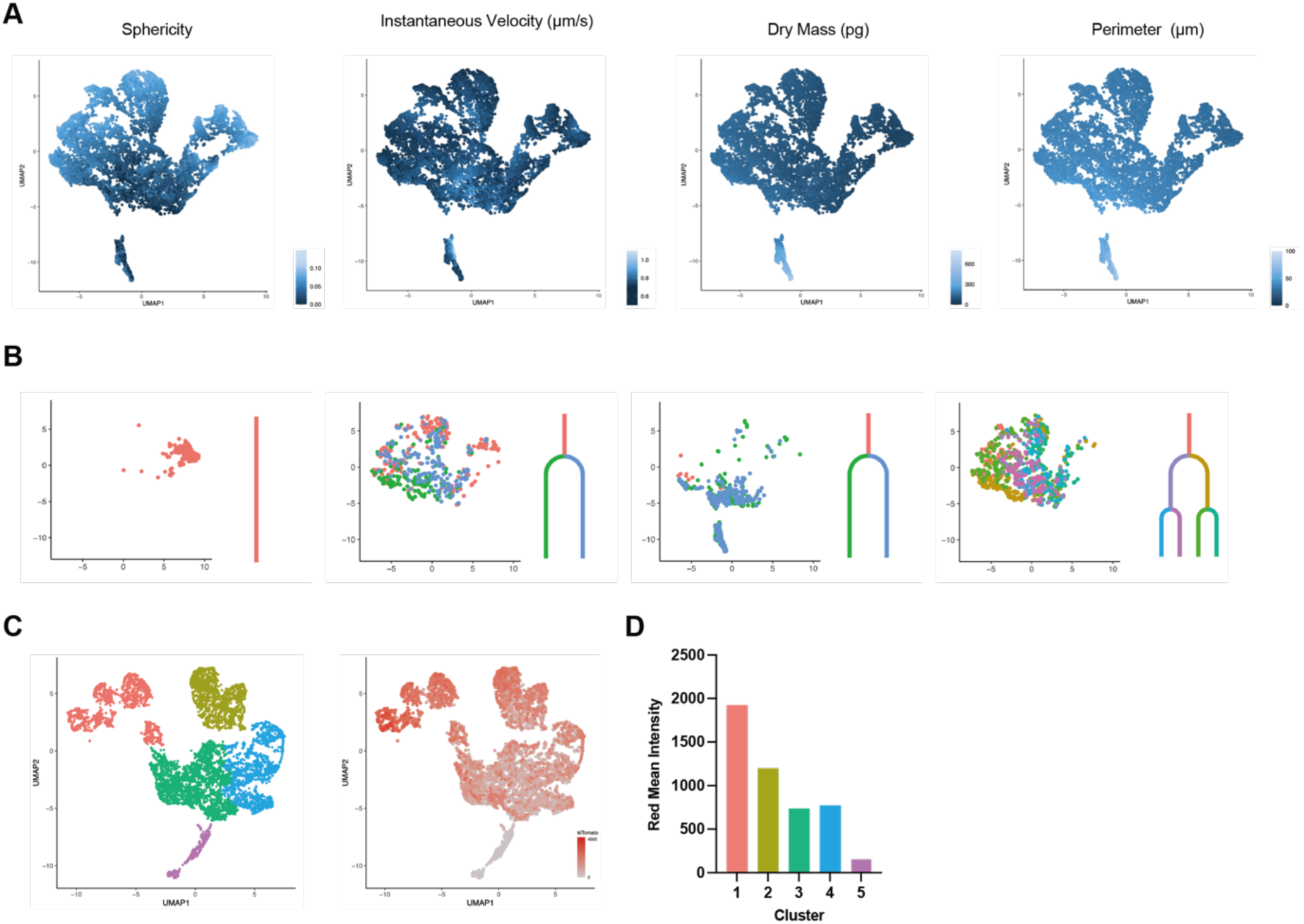
Prediction of *Hlf* expression levels based on cellular kinetic features. **(A**) UMAP representation overlaying sphericity, velocity, dry mass and perimeter. **(B)** Representative cell tracking on the UMAP plot. Each color represents a distinct division. **(C)** UMAP plot colored by hierarchical clustering using a different dataset (left) and mean red fluorescence intensity per cluster (right). **(D)** Mean red fluorescence intensity of Hlf-tdTomato across clusters.

**Fig. S5.**
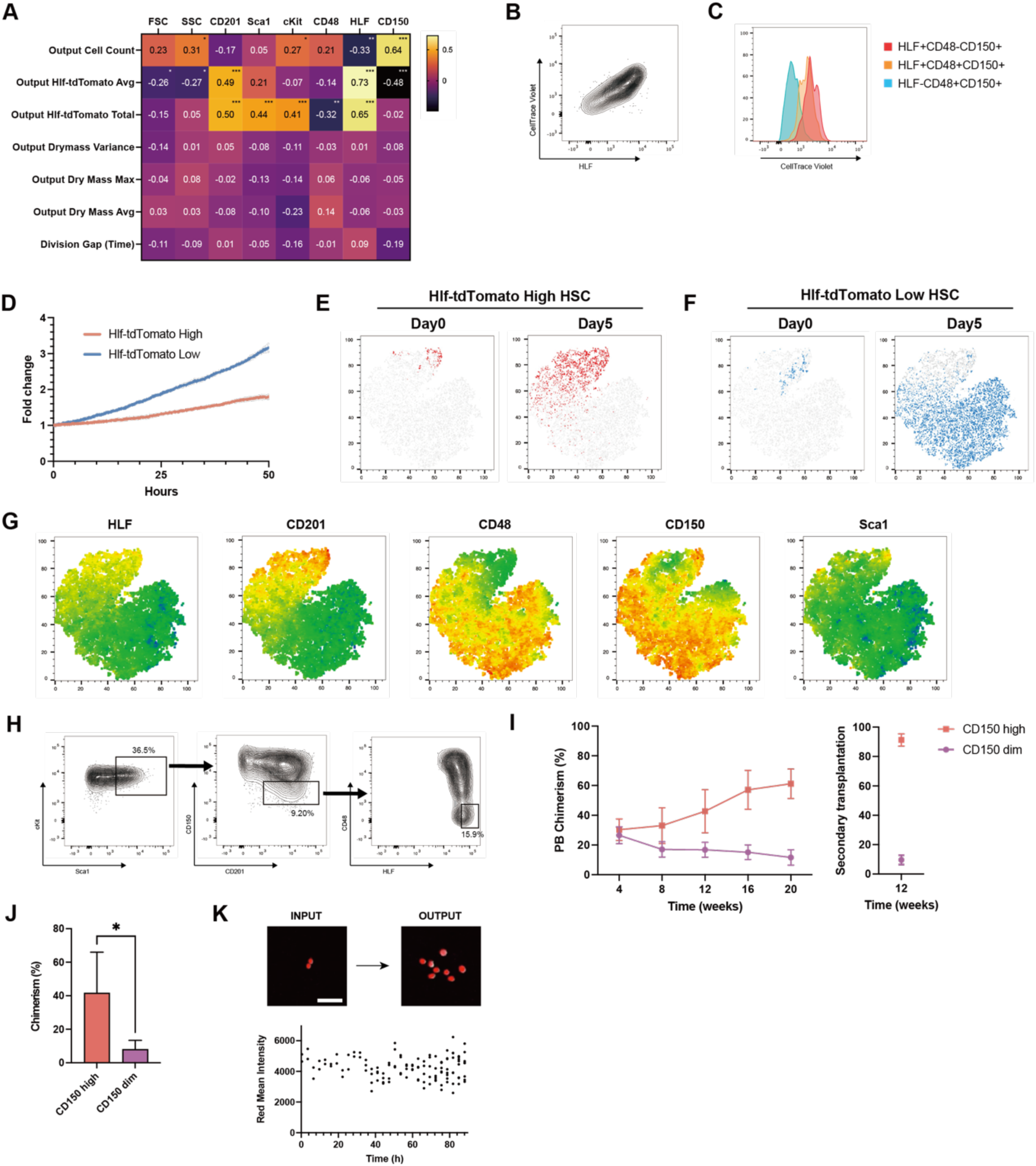
Cellular dynamics of expanded Hlf-tdTomato HSCs. **(A**) Heatmap showing the correlation between index data and cellular dynamics. Statistical significance was calculated using a t-test. *P<0.05, **P<0.01, ***P<0.001. **(B, C)** Analysis of the relationship between HSC division frequency and Hlf-tdTomato levels using CellTrace. A representative plot shows the correlation between CellTrace Violet and Hlf-tdTomato (B), and immature HSPC fractions show higher CellTrace Violet levels. **(D)** Cellular proliferation dynamics of Hlf-tdTomato high/low expanded HSCs. Fifty HSCs were expanded for 50 hours and monitored by QPI. **(E-G)** tSNE plots of expanded HSCs based on FACS data using CD201, CD48, CD150, cKit, Sca1, and Lin markers. Complete data from day 0 high and low Hlf-tdTomato HSCs and after 5 days of expansion were combined for tSNE analysis. High Hlf-tdTomato HSCs at day 0 and after 5 days are shown in red (E), while low Hlf-tdTomato HSCs at day 0 and after 5 days are shown in blue (F). tSNE plots are colored by Hlf-tdTomato, CD201, CD48, CD150, and Sca1 (G). **(H)** Gating strategy for CD150^dim^ Hlf-tdTomato high CD201^+^cKit^+^Sca1^+^Lin^-^ HSCs. **(I)** Mean donor peripheral blood chimerism in primary recipients (n = 4, 5 mice per group) and secondary recipients (n = 3–5 mice per group). **(J)** Mean donor bone marrow chimerism in primary recipients (n = 4, 5 mice per group). Error bars represent SD. Statistical significance was calculated using a t-test. *P<0.05 **(K)** Representative images of HSCs that exceptionally maintained high levels of Hlf-tdTomato and produced a large number of cells.

## SUPPLEMENTAL MOVIE TITLES AND LEGENDS

**Movie S1**

Representative videos of quantitative phase imaging during a single-HSC ex vivo expansion. The left video shows cells that divide rapidly and produce a large number of cells, while the right video shows cells that divide slowly and produce only a small number of cells.

**Movie S2**

Representative videos of quantitative phase imaging during a single-HSC ex vivo expansion. The left video shows cells that produce cells with high dry mass, while the right video shows cells that produce cells with low dry mass.

**Movie S3**

Representative videos of quantitative phase imaging during a single-HSC ex vivo expansion. The left video shows cells with “Division Gap” of less than 5 hours, where the second division occurs almost simultaneously. The right video shows cells with “Division Gap” of more than 5 hours.

**Movie S4**

Representative videos of quantitative phase imaging of bulk expanded HSCs over a wide field of 1000 µm. HSCs were expanded for one week, followed by 36 hours of imaging. The latter part of the video demonstrates an example of cell division detection using Trackmate.

**Movie S5**

Representative video of quantitative phase imaging of bulk expanded HSCs in a U-bottom plate.

**Movie S6**

Representative video of quantitative phase imaging and fluorescence imaging of bulk expanded Hlf-tdTomato HSCs in a U-bottom plate.

**Movie S7**

Representative videos of quantitative phase imaging and fluorescence imaging during a single-HSC ex vivo expansion. The left video shows cells that divide slowly maintaining a high level of tdTomato expression, whereas the right video shows cells that divide quickly and exhibit a decrease in tdTomato expression levels.

**Movie S8**

Representative video of quantitative phase imaging and fluorescence imaging during a single-HSC ex vivo expansion. The video shows cells that divide maintaining high levels of tdTomato expression but undergo early cell death, resulting in no increase in the total cell count.

**Movie S9**

Representative video of quantitative phase imaging and fluorescence imaging during a single-HSC ex vivo expansion. The video shows cells that actively divide while exceptionally maintaining high levels of tdTomato expression.

**Movie S10**

Representative videos of quantitative phase imaging of expanded HSCs with and without cell motility. The left video shows cells with movement, while the right video shows cells fixed in the center, with motility information removed.

## References

1. J. E. Till, E. A. McCulloch, A direct measurement of the radiation sensitivity of normal mouse bone marrow cells. Radiat. Res. 14, 213–222 (1961).

2. J. W. M. Visserc, J. G. J. Bauman, A. H. Mulder, J. F. Euason, A. M. De Leeuw, Isolation of murine pluripotent hemopoietic stem cells. J. Exp. Med. 159, 1576–1590 (1984).

3. G. J. Spangrude, S. Heimfeld, I. L. Weissman, Purification and characterization of mouse hematopoietic stem cells. Science (80-.). 241, 58–62 (1988).

4. S. J. Morrison, I. L. Weissman, The long-term repopulating subset of hematopoietic stem cells is deterministic and isolatable by phenotype. Immunity. 1, 661–673 (1994).

5. M. Osawa, K. I. Hanada, H. Hamada, H. Nakauchi, Long-term lymphohematopoietic reconstitution by a single CD34-low/negative hematopoietic stem cell. Science (80-.). 273, 242–245 (1996).

6. F. Notta, S. Doulatov, E. Laurenti, A. Poeppl, I. Jurisica, J. E. Dick, Isolation of single human hematopoietic stem cells capable of long-term multilineage engraftment. Science (80-.). 333, 218–221 (2011).

7. J. Y. Chen, M. Miyanishi, S. K. Wang, S. Yamazaki, R. Sinha, K. S. Kao, J. Seita, D. Sahoo, H. Nakauchi, I. L. Weissman, Hoxb5 marks long-term haematopoietic stem cells and reveals a homogenous perivascular niche. Nat. 2016 5307589. 530, 223 (2016).

8. Y. Tajima, K. Ito, A. Umino, A. C. Wilkinson, H. Nakauchi, S. Yamazaki, Continuous cell supply from Krt7-expressing hematopoietic stem cells during native hematopoiesis revealed by targeted in vivo gene transfer method. Sci. Reports 2017 71. 7, 1–10 (2017).

9. K. Ito, S. Yamazaki, R. Yamamoto, Y. Tajima, A. Yanagida, T. Kobayashi, M. Kato-Itoh, S. Kakuta, Y. Iwakura, H. Nakauchi, A. Kamiya, Gene Targeting Study Reveals Unexpected Expression of Brain-expressed X-linked 2 in Endocrine and Tissue Stem/Progenitor Cells in Mice. J. Biol. Chem. 289, 29892–29911 (2014).

10. M. A. Goodell, H. Nguyen, N. Shroyer, Somatic stem cell heterogeneity: diversity in the blood, skin and intestinal stem cell compartments. Nat. Rev. Mol. Cell Biol. 2015 165. 16, 299–309 (2015).

11. L. E. Purton, Adult murine hematopoietic stem cells and progenitors: an update on their identities, functions, and assays. Exp. Hematol. 116, 1–14 (2022).

12. B. Rix, A. H. Maduro, K. S. Bridge, W. Grey, Markers for human haematopoietic stem cells: The disconnect between an identification marker and its function. Front. Physiol. 13, 1009160 (2022).

13. E. Laurenti, B. Göttgens, From haematopoietic stem cells to complex differentiation landscapes. Nature. 553 (2018), pp. 418–426.

14. G. Donati, F. M. Watt, Stem Cell Heterogeneity and Plasticity in Epithelia. Cell Stem Cell. 16, 465–476 (2015).

15. J. Carrelha, Y. Meng, L. M. Kettyle, T. C. Luis, R. Norfo, V. Alcolea, H. Boukarabila, F. Grasso, A. Gambardella, A. Grover, K. Högstrand, A. M. Lord, A. Sanjuan-Pla, P. S. Woll, C. Nerlov, S. E. W. Jacobsen, Hierarchically related lineage-restricted fates of multipotent haematopoietic stem cells. Nat. 2018 5547690. 554, 106–111 (2018).

16. A. Sanjuan-Pla, I. C. Macaulay, C. T. Jensen, P. S. Woll, T. C. Luis, A. Mead, S. Moore, C. Carella, S. Matsuoka, T. B. Jones, O. Chowdhury, L. Stenson, M. Lutteropp, J. C. A. Green, R. Facchini, H. Boukarabila, A. Grover, A. Gambardella, S. Thongjuea, J. Carrelha, P. Tarrant, D. Atkinson, S. A. Clark, C. Nerlov, S. E. W. Jacobsen, Platelet-biased stem cells reside at the apex of the haematopoietic stem-cell hierarchy. Nat. 2013 5027470. 502, 232–236 (2013).

17. A. E. Rodriguez-Fraticelli, S. L. Wolock, C. S. Weinreb, R. Panero, S. H. Patel, M. Jankovic, J. Sun, R. A. Calogero, A. M. Klein, F. D. Camargo, Clonal analysis of lineage fate in native haematopoiesis. Nature. 553, 212–216 (2018).

18. R. Yamamoto, Y. Morita, J. Ooehara, S. Hamanaka, M. Onodera, K. L. Rudolph, H. Ema, H. Nakauchi, Clonal Analysis Unveils Self-Renewing Lineage-Restricted Progenitors Generated Directly from Hematopoietic Stem Cells. Cell. 154, 1112–1126 (2013).

19. E. Mansell, V. Sigurdsson, E. Deltcheva, J. Brown, C. James, K. Miharada, S. Soneji, J. Larsson, T. Enver, Mitochondrial Potentiation Ameliorates Age-Related Heterogeneity in Hematopoietic Stem Cell Function. Cell Stem Cell. 28, 241–256.e6 (2021).

20. N. Vannini, M. Girotra, O. Naveiras, G. Nikitin, V. Campos, S. Giger, A. Roch, J. Auwerx, M. P. Lutolf, Specification of haematopoietic stem cell fate via modulation of mitochondrial activity. Nat. Commun. 2016 71. 7, 1–9 (2016).

21. H. Zhou, I. Li, C. S. Bramlett, B. Wang, J. Hao, D. P. Yen, Y. Ando, S. E. Fraser, R. Lu, K. Shen, Label-free metabolic optical biomarkers track stem cell fate transition in real time. Sci. Adv. 10, 6770 (2024).

22. S. Nestorowa, F. K. Hamey, B. Pijuan Sala, E. Diamanti, M. Shepherd, E. Laurenti, N. K. Wilson, D. G. Kent, B. Göttgens, A single-cell resolution map of mouse hematopoietic stem and progenitor cell differentiation. Blood. 128, e20–e31 (2016).

23. Di. Loeffler, T. Schroeder, Understanding cell fate control by continuous single-cell quantification. Blood. 133, 1406–1414 (2019).

24. T. Kull, T. Schroeder, Analyzing signaling activity and function in hematopoietic cells. J. Exp. Med. 218 (2021), doi:10.1084/JEM.20201546/212378.

25. A. C. Wilkinson, R. Ishida, M. Kikuchi, K. Sudo, M. Morita, R. V. Crisostomo, R. Yamamoto, K. M. Loh, Y. Nakamura, M. Watanabe, H. Nakauchi, S. Yamazaki, Long-term ex vivo haematopoietic-stem-cell expansion allows nonconditioned transplantation. Nature. 571, 117–121 (2019).

26. H. J. Becker, R. Ishida, A. C. Wilkinson, T. Kimura, M. S. J. Lee, C. Coban, Y. Ota, Y. Tanaka, M. Roskamp, T. Sano, A. Tojo, D. G. Kent, S. Yamazaki, Controlling genetic heterogeneity in gene-edited hematopoietic stem cells by single-cell expansion. Cell Stem Cell. 30, 987–1000.e8 (2023).

27. M. Sakurai, K. Ishitsuka, R. Ito, A. C. Wilkinson, T. Kimura, E. Mizutani, H. Nishikii, K. Sudo, H. J. Becker, H. Takemoto, T. Sano, K. Kataoka, S. Takahashi, Y. Nakamura, D. G. Kent, A. Iwama, S. Chiba, S. Okamoto, H. Nakauchi, S. Yamazaki, Chemically defined cytokine-free expansion of human haematopoietic stem cells. Nat. 2023 6157950. 615, 127–133 (2023).

28. J. M. Rodenburg, M. J. Humphry, A. M. Maiden, Optical ptychography: a practical implementation with useful resolution. Opt. Lett*. Vol.* 35*, Issue* *15*, pp. 2585-2587. 35, 2585–2587 (2010).

29. A. M. Maiden, J. M. Rodenburg, An improved ptychographical phase retrieval algorithm for diffractive imaging. Ultramicroscopy. 109, 1256–1262 (2009).

30. T. Yokomizo, N. Watanabe, T. Umemoto, J. Matsuo, R. Harai, Y. Kihara, E. Nakamura, N. Tada, T. Sato, T. Takaku, A. Shimono, H. Takizawa, N. Nakagata, S. Mori, M. Kurokawa, D. G. Tenen, M. Osato, T. Suda, N. Komatsu, Hlf marks the developmental pathway for hematopoietic stem cells but not for erythro-myeloid progenitors. J. Exp. Med. 216, 1599–1614 (2019).

